# Myelin-informed forward models for M/EEG source reconstruction

**DOI:** 10.1101/2024.06.30.601378

**Authors:** S Helbling, SS Meyer, N Weiskopf

## Abstract

Magnetoencephalography (MEG) and Electroencephalography (EEG) provide direct electrophysiological measures at an excellent temporal resolution, but the spatial resolution of source-reconstructed current activity is limited to several millimetres. Here we show, using simulations of MEG signals and Bayesian model comparison, that non-invasive myelin estimates from high-resolution quantitative magnetic resonance imaging (MRI) can enhance MEG/EEG source reconstruction. Our approach assumes that MEG/EEG signals primarily arise from the synchronised activity of pyramidal cells, and since most of the myelin in the cortical sheet originates from these cells, myelin density can predict the strength of cortical sources measured by MEG/EEG. Leveraging recent advances in quantitative MRI, we exploit this structure-function relationship and scale the leadfields of the forward model according to the local myelin density estimates from in vivo quantitative MRI to inform MEG/EEG source reconstruction. Using Bayesian model comparison and dipole localisation errors (DLEs), we demonstrate that adapting local forward fields to reflect increased local myelination at the site of a simulated source explains the simulated data better than models without such leadfield scaling. Our model comparison framework proves sensitive to myelin changes in simulations with exact coregistration and moderate-to-high sensor-level signal-to-noise ratios (≥10 dB) for the multiple sparse priors (MSP) and empirical Bayesian beamformer (EBB) approaches. Furthermore, we sought to infer the microstructure giving rise to specific functional activation patterns by comparing the myelin-informed model which was used to generate the activation with a set of test forward models incorporating different myelination patterns. We found that the direction of myelin changes, however not their magnitude, can be inferred by Bayesian model comparison. Finally, we apply myelin-informed forward models to MEG data from a visuo-motor experiment. We demonstrate improved source reconstruction accuracy using myelin estimates from a quantitative longitudinal relaxation (R1) map and discuss the limitations of our approach.

**Highlights:**

- We use quantitative MRI to implement myelin-informed forward models for M/EEG
- Local myelin density was modelled by adapting the local leadfields
- Myelin-informed forward models can improve source reconstruction accuracy
- We can infer the directionality of myelination patterns, but not their strength
- We apply our approach to MEG data from a visuo-motor experiment

## Introduction

Magnetoencephalography (MEG) and EEG (Electroencephalography) provide direct electrophysiological measures of synchronous neural activity primarily involving cortical pyramidal cells at an excellent temporal resolution. However, estimating the spatial distribution of neural sources in the brain from MEG/EEG sensor data outside the head is an ill-posed inverse problem. It is widely recognised that inverse solutions of MEG/EEG data for cortical activity are not unique^1^.

To address the ill-posed nature of the inverse problem, prior assumptions on the distribution of neural sources must be incorporated. Various assumptions regarding the current source distribution can be implemented by introducing priors on the source covariance. For instance, these priors can specify whether the current sources are expected to be uniformly distributed across the cortical surface (Minimum Norm Estimates; MNE^2^) or exhibit sparsity (Multiple Sparse Priors; MSP^3^). Additionally, anatomical priors can be applied, e.g., by restricting the source space to the cortical surface and constraining sources to be oriented perpendicular to the local cortical surface^4^. Anatomical information has also been incorporated to differentiate between current sources originating from deep or superficial laminae or to distinguish hippocampal from cortical sources^5–9^.

In this study, we propose a novel anatomical approach that uses myelin estimates derived from high-resolution quantitative MRI (qMRI) to enhance MEG/EEG source reconstruction accuracy. Quantitative MRI combines traditional ‘weighted’ MR images in a model-based manner to produce reproducible and standardised measures in physical units, which are less dependent on the acquisition scheme^10–14^. Crucially, qMRI is sensitive to microstructural properties of brain tissue such as axon, myelin, iron and water concentration^14–20^ and thus presents an opportunity to move beyond classical histology to qMRI-based *in vivo* characterisation of brain microstructure^10,14^. Of particular interest for our study, several qMRI maps enable non-invasive estimation of myelin content throughout the cortex^21–23^. Specifically, we will focus on longitudinal relaxation rate (R1) maps from a multi-parameter map protocol^24^ when applying our approach to experimental data. R1 maps have been validated as a marker for myelin^20,25,26^, showing high test-retest reliability and good inter-site reliability^24,27,28^.

While myelin is important for fast propagation of action potentials, axon protection, trophic support and learning^29–32^, our focus here is on using myelin as a proxy for cell density, as we assume that cell density is predictive of MEG signal strength. Myelin-sensitive qMRI measures in the cortex are expected to correlate positively with local cell density due to the close relationship between cyto- and myeloarchitecture^33–35^. Direct links between qMRI and cytoarchitecture have been further demonstrated by positive correlations between R2* measures at 7T and cell counts from the von Economo atlas^36^, as well as cell type-specific gene enrichment analyses showing significant associations between R1 and R2* maps with genes enriched in GABA- and glutamatergic neurons^37^. As M/EEG signals are predominantly generated by cortical pyramidal cells, we expect qMRI measures to predict MEG signal strength. Importantly, we have previously demonstrated a positive correlation between MEG responses and cortical grey matter myelin estimates from quantitative MRI across participants^38^. This association renders myelin estimates from qMRI useful as a structural prior to improve MEG source reconstruction accuracy.

Here we propose a novel approach that uses this myelin information as a histological constraint to improve M/EEG source reconstruction. This is done by scaling the leadfields of the forward model based on the qMRI-derived local myelination. More precisely, we increase the leadfield strength at areas with higher local myelination and decrease it in areas with lower local myelination. We thus introduce a prior at the level of the forward model that reflects our expectation of a higher current source density at more strongly myelinated cortical regions.

The outline of this paper is the following: We introduce a novel approach to improve M/EEG source reconstruction using myelin-informed forward models and aim to investigate the conditions under which myelin-informed generative models can enhance M/EEG source reconstruction. We systematically test different source reconstruction priors, signal-to-noise ratios (SNRs), myelin scaling factors and co-registration errors to determine the impact of these factors. Additionally, we assess the potential of comparing different myelin-informed generative models to infer the underlying microstructure. We then apply our approach to experimental data from a visuo-motor paradigm and demonstrate that myelin-informed generative models have the potential to improve the accuracy of M/EEG source reconstructions for experimental MEG data.

## Methods

We conduct simulations of MEG activity to investigate whether integrating MRI-based histology into MEG generative models can improve source reconstruction accuracy. To this aim, we adapted the leadfields of the MEG forward model to mimic changes in local myeloarchitecture and evaluated dipole localisation errors (DLEs) and Bayesian model evidence of the source reconstructed data in dependence of myelination patterns. When using Bayesian model evidence, our guiding assumption is that incorporating true structural information into the forward model would result in lower model complexity and, consequently, lower model evidence^5,8,39–41^.

Simulations and data analyses were implemented using the SPM12 software package (http://www.fil.ion.ucl.ac.uk/spm/software/spm12/) and in-house code in Matlab (The Mathworks Inc., Natick, MA, USA). The code necessary to replicate the simulations presented in this paper is available via GitHub (http://github.com/sashel/myelin_sim)

### Single dipole simulations

We first ask whether modelling increased myelination at the location of a simulated source by scaling the local leadfields can improve source reconstruction accuracy. This was tested across a range of popular source reconstruction approaches, signal-to-noise ratios (SNRs), and levels of co-registration errors to determine the constraints under which myelin-informed forward models lead to a significant improvement in source reconstruction accuracy.

We based our simulations on a single dataset acquired with a CTF 275-channel system. This experimental MEG dataset was used to define head positions and channel locations, while structural MRIs of the same participant (see subsection “MRI data acquisition”) were used to define the forward model. Synthetic datasets with single patches of activated cortex were generated by simulating a sinusoidal dipolar source of 20 Hz for 300 ms (6 cycles) with a total effective dipole moment of 20 nAm, similar to previous simulation studies^5,8^. Each simulated single-trial dataset had a duration of 800 ms, epoched from -200 to 600 ms, with the dipolar source modelled between 100 and 400 ms.

We used 60 locations based on 30 out of 50 randomly selected bilateral patch pairs as cortical source locations. Gaussian source patches are defined within SPM12 as follows^42^:

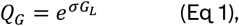

where *σ* determines the spatial extent of the activated patch and *G*_*L*_ denotes the graph Laplacian 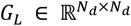, with *N*_*d*_ current dipoles distributed through the cortical surface.

The graph Laplacian is based on an adjacency matrix 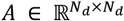, with *A*_*ij*_ = 1 if there is face connectivity, and zero otherwise, and is defined as:

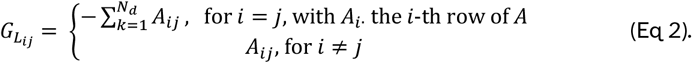

The full width at half maximum of each simulated patch was set to 6 mm, and patches spanned on average 14.0667 ± 3.349 vertices.

We simulated single trial data of averaged evoked activity instead of datasets with multiple trials to reduce the computational load of our simulations. Random Gaussian white noise was added to the sensor level data at different SNRs (0, -5, -10, -15, -20 dB) for each of these single trial datasets.

### Source Reconstruction

The leadfield matrix describing the sensitivity of the MEG sensors to the cortical sources was computed using the Nolte Single Shell forward model^43^. We generated 60 synthetic datasets, one for each dipole location, with standard, non-adapted leadfields. For source reconstruction, we scaled the leadfields to mimic increased or decreased myelination at the location of the simulated dipole patch. These adapted forward models reflect our expectation that regions of higher myelination will have higher cortical current densities by weighting the leadfields accordingly. Scaling factors were 5, 3, and 3/2, and their reciprocals 1/5, 1/3, and 2/3 to simulate increases or decreases in myelination compared to the overall myelination across the cortex. Note that we applied a weighting such that the leadfield adaptation followed the Gaussian shape of the simulated dipole moment, i.e., the leadfield scaling was further weighted by the simulated signal magnitude at each vertex with the strongest scaling occurring at the peak vertex:

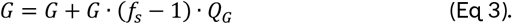

With an average patch weighting of about 0.2 at the peak vertex of patches, a myelin scaling factor of 5 translates to an adapted leadfield that is 1.8 times larger than the original leadfield.

Source reconstruction for each simulated dataset was applied to a Hanning windowed time window of 300 ms, covering the whole duration of the simulated cortical activation from 100 to 400 ms. Data were low-pass filtered at 80 Hz before inversion. Source inversion was performed using Bayesian source reconstruction with three popular priors within SPM12: minimum norm estimate^2^ (MNE), empirical Bayesian beamformer^42,44^ (EBB) and multiple sparse priors (MSP^3^) to estimate the underlying current sources from the simulated sensor data. We used the classic Bayesian source inversion scheme in SPM12 without any of the re-scaling factors that are implemented in SPM to allow for mixing of different imaging modalities or group imaging^45^.

Bayesian model comparison was used to determine if adapting the forward model to incorporate information on cortical myelination led to improved source reconstruction accuracy compared to a standard forward model. Log model evidence across source reconstruction solutions based on the different myelin-informed and standard forward models was approximated using Variational Free energy. Free energy is a parametric metric that rewards accuracy and penalises model complexity^42,46^, providing a lower bound for the log model evidence value^47^. A difference in log model evidence greater than 3 suggests that one model is approximately 20 times more likely than another.

Dipole localization errors were also used as a metric to investigate source reconstruction accuracy and were defined as the Euclidean distance between the true source location of the simulated activation and the maximum of the estimated source distribution. Significance of DLE differences were evaluated for each source reconstruction approach, SNR and scaling factor by comparing the DLEs obtained for the myelin-informed forward model with the DLEs of the standard null model without leadfield scaling using a paired t-test.

To evaluate the spatial specificity of myelin-informed generative models, we ran an additional set of control simulations where we simulated single patches of activated cortex at the same 60 locations but adapted the leadfields at the patch location at the opposite hemisphere.

In addition, to investigate the impact of co-registration errors, we ran a further set of simulations where we added random errors of 1 to 5 mm in 1 mm increments to each of the three fiducial locations before inverting the model. This changes the affine transformation matrix that aligns the fiducials of the MEG sensor array and the structural MRI and hence the cortical source locations that are based on the MRI-derived cortical surface relative to the MEG sensors. For each dataset with added random co-registration error, we thus calculated a new gain matrix, whose leadfields were then adapted and used for source reconstruction.

### Inferring underlying myelination patterns

In a next set of simulations, we aimed to infer the underlying myelination pattern from the simulated sensor activity alone. To model myelination patterns, we simulated cortical activity at both patch locations of the 30 bilateral pairs simultaneously, increasing the leadfield strength at one and decreasing it at the opposite location. Unlike as in the single dipole simulations where leadfields were only adapted at the level of source reconstruction, when modelling myelination patterns, we adapted the leadfields prior to the generation of sensor data, i.e., source locations with increased (decreased) leadfield strength would have a stronger (weaker) impact on the sensor level data. We combined the six scaling factors used for the single dipole simulations into scaling pairs to construct six myelin-informed forward models with different scaling strengths and directions (5 – 1/5,3 – 1/3, 3/2 – 2/3 as well as 1/5 – 5, 1/3 – 3 and 2/3 – 3/2), along with a null model without leadfield modifications (1 – 1). Each dataset was then source reconstructed using the EBB approach and variational free energy was obtained across the same set of standard and myelin-informed forward models. Bayesian model comparison was used to select the best model based on these log model evidence estimates^46^.

As correlated sources can lead to the reduction of the estimated source strength when using beamformers, we used bandlimited white noise waveforms between 1-80 Hz for 300 ms instead of sinusoidal sources for the paired source patches in the myelination pattern simulations. The effective dipole moments were set to 20 nAm, as in the single dipole simulations. All myelination pattern simulations were performed without added coregistration errors.

### Application to experimental MEG data from a visuo-motor paradigm

We next applied myelin-informed generative models to MEG data from a visuo-motor experiment to demonstrate that our approach can improve source reconstruction accuracy for experimentally measured MEG data. The MEG data were acquired from 14 participants (mean age 27.2 years, std: 4.0 y, 5 females, 9 males) as part of the MEG UK Normative database project at WCNC, UCL, London. In addition, quantitative MRIs from a multi-parameter maps protocol were acquired for each participant as part of the project. In each trial of the experiment, a vertical, stationary, square-wave grating with a frequency of 3 cycles per degree and covering approximately 4 × 4° on a mean luminance grey background was shown for 1.5 to 2 seconds in the lower visual field of the left or the right hemisphere, followed randomly by a short or a long inter-stimulus interval (ISI, 4 or 8 sec). Grating displays were generated using the Psychophysics Toolbox^48^ in Matlab. Participants were instructed to fixate a central white cross while covertly attending the gratings and to perform an abduction movement with their left index finger each time the grating stimulus disappeared. We presented 200 trials in two blocks of approximately 13 min duration each, with the presentation site randomised across trials. The study was approved by the University College London ethics committee (reference number 3090/001), and written informed consent was obtained from all participants prior to scanning.

#### MEG data acquisition and preprocessing

The MEG data acquisition and preprocessing were performed using a 275-channel whole-head MEG setup with synthetic third-order gradiometers (CTF systems) at a 1200 Hz sampling rate. The MEG was located inside a magnetically shielded room to minimise external electromagnetic noise. Subjects wore custom-made 3D-printed headcasts during scanning to minimise head movements and increase co-registration accuracy, as described in previous studies^49,50^. Head location was continuously determined during each run using head coils at three fiducial positions, the nasion and left and right preauricular points. Head coils were placed inside indentations of the headcasts for highly accurate co-registration with the structural MRI. An electromyogram (EMG) was recorded using three electrodes at the left hand to determine the onset of finger abduction movements. Data were converted for analysis in SPM12, downsampled to 300 Hz and epoched into trials of 3.2 seconds ([-0.60 2.60] sec with respect to the grating onset). We analysed only correct responses in which participants responded with a finger abduction movement between 100 and 900 ms after the grating offset. Any epochs containing peak-to-peak amplitude signals greater than 5×10^−12^ T were classified as artefacts and removed from the analysis. Independent component analysis (ICA) using the Extended Infomax algorithm was applied to the concatenated MEG data of the remaining epochs to remove eye movement and heartbeat artefacts.

#### MRI data acquisition

Prior to the MEG measurement, participants underwent two MRI scanning protocols during the same visit. Subjects were scanned on a 3T whole body MR system (Magnetom TIM Trio, Siemens Healthcare, Erlangen, Germany). The first protocol was used to generate an accurate image of the scalp for headcast construction as described in Meyer et al., 2017. Special care was taken to prevent distortions in the image due to skin displacement on the face, head, or neck, as any such errors could compromise the fit of the headcast. Accordingly, a more spacious 12 channel head coil than the standard 32 channel head coil was used for the headcast scan. Acquisition time was 3 min 42 sec.

In a second protocol, multi-parameter mapping was performed using spoiled multi-echo 3D fast low angle shot (FLASH) acquisitions with predominantly proton density (PD), T1 or magnetization transfer (MT) weighting according to the MPM protocol^24^. A 32-channel head coil was used to increase SNR. The MPM data were acquired with whole-brain coverage at an isotropic resolution of 800 μm using a FoV of 256 mm (H-F), 224 mm (A-P), and 179 mm (R-L). The flip angle was 6 degrees for the PD- and MT-weighted volumes and 21 degrees for the T1-weighted acquisition. MT-weighting was achieved through the application of a Gaussian RF pulse 2kHz off resonance with 4 ms duration and a nominal flip angle of 220°.

Gradient echoes were acquired with alternating readout gradient polarity at eight equidistant echo times ranging from 2.34 to 18.44 ms in steps of 2.30 ms using a readout bandwidth of 488 Hz/pixel. Only six echoes were acquired for the MT-weighted acquisition to incorporate the RF pulse and to maintain a repetition time (TR) of 25 ms for all FLASH volumes. To accelerate the data acquisition, partial parallel imaging using the GRAPPA algorithm was employed in each phase-encoded direction (AP and RL) with a speed-up factor of two.

To maximise the accuracy of the measurements, inhomogeneity in the RF transmit field was mapped using a 3D echo (EPI) acquisition of spin and stimulated echoes with 15 different refocusing flip angles (TE/TM/TR = 39.38/33.24/500 ms; matrix = 64 × 48 × 48 pixels; FoV = 256 × 192 × 192 mm) following the approach described in ^51^. A B0 field-map was acquired using a double-echo gradient echo sequence (TE1 = 10 ms, TE2 = 12.46 ms, TR=1020 ms, 3 × 3 × 2 mm resolution, 1 mm gap; matrix size = 64 × 64 pixels; FoV = 192 × 192 × 191 mm) to allow for post-processing correction of geometric distortions of the EPI data due to B0 field inhomogeneity. Total acquisition time for all MRI scans was less than 30 min.

#### Estimation of qMRI parameter maps

Quantitative maps were calculated using the hMRI toolbox for quantitative MRI and in vivo histology using MRI^52^ within the SPM12 framework. Maps of the effective transverse relaxation rate R2* were estimated from the gradient echoes of all contrasts using the ordinary least squares ESTATICS approach^53^. The proton-density weighted (PDw), T1-weighted (T1w), and magnetisation transfer-weighted (MTw) data were averaged over the first six echoes to increase the SNR^54^, and the three resulting volumes were used to calculate MT, R1, and effective proton density (PD*) maps as previously described^24,55^. For myelin mapping, we focused on the R1 maps. The effective proton density map PD* was also used, together with the R1 map, for cortical surface reconstruction within the Freesurfer pipeline.

#### Cortical surface reconstruction

We run a bespoken Freesurfer^56^ (version 6.0.0) pipeline tailored to MPM-based input images to generate cortical surfaces for the white matter - grey matter (“white” surface) and the grey matter-pial (“pial” surface) boundaries. First, a small number of negative and very high values were pruned to 0 in the R1 and PD maps using AFNI, such that *T*1 (= 1/*R*1) was bounded between [0, 8000] ms and PD between [0, 150] %. Then, the PD and *T*1 maps were used as input to FreeSurfer’s mri_synthesize routine to create a synthetic FLASH volume with optimal white matter (WM)/grey matter (GM) contrast (TR 20 ms, FA 30°, TE 2.5 ms).

This synthesised image was provided as input to the initial part of the Freesurfer pipeline, autorecon-1, but with the skull-stripping flag set to -no. Skull stripping was performed using SPM_segment to construct a combined GM/WM/cerebrospinal fluid (CSF) brain mask (threshold for each tissue class: tissue probability > 0). The skull-stripped image was then used as input for the remaining steps of the recon-all pipeline to reconstruct cortical surfaces.

A mid-cortical surface was generated by expanding the white matter surface to 50% of the local cortical depth for each hemisphere. Mid-cortical surfaces of both hemispheres were aligned to the headcast scan, converted to GIfTI file format, and combined into a single surface for each participant. Each of these combined surfaces was downsampled by a factor of 10 using Freesurfer’s mris_decimate function, resulting in cortical surface meshes with on average 29.536 vertices.

#### Myelin-informed M/EEG forward models visuo-motor paradigm

The leadfields of the forward model were computed using the Nolte corrected-sphere approach^43^ with a template-derived canonical surface for the inner skull surface^57^ and a subject-specific mid-cortical surface that defined the source space. Leadfields were oriented normally to the cortical surface mesh. Note that while we constructed the cortical surfaces from the MPM scans, both the MPMs and their cortical surfaces were transformed to headcast space via co-registration to the headcast scan before calculating the forward model. This approach was chosen to exploit the excellent co-registration accuracy between MEG sensors and the headcast scan, which is due to the precise knowledge of the fiducial locations for the latter.

For the experimental data, myelin-informed forward models were constructed by scaling the leadfields by the myelin density values estimated from the R1 maps. Myelin estimates at 50% cortical depth were smoothed across the cortical surface with a full-width half maximum (FWHM) of 3 mm using Freesurfer. Subsequently, myelin estimates at a given vertex were normalised by subtracting the mean and dividing by the standard deviation of all myelin estimates sampled across the cortical surface. Each leadfield was then scaled by the product of these normalised myelin estimates *f*_*norm*_ with a scaling factor *f*_*s*_ that ranged from 0.02, 0.05, 0.2 to 5 resp. -0.02, -0.05, -0.2 to -5 in the following way:

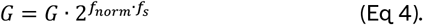

Scaling factors *f*_*s*_ could be positive or negative, corresponding to the hypotheses that increased myelination leads to a weaker or stronger current source density, respectively. As an example, at a scaling factor *f*_*s*_ of 0.2, a normalised myelin estimate *f*_*norm*_ of 2 at a given vertex translates to a leadfield that is approximately 132% as large as the original one, while at a negative scaling factor *f*_*s*_ of -0.2 the same *f*_*norm*_ translates to a leadfield of about 76% of the strength of the original leadfield.

As the mid-cortical layer is not well-defined in non-isocortical areas, non-isocortical areas were omitted from the scaling procedure and leadfields at those areas were not adapted by myelin density estimates. Non-isocortical areas included retrosplenial Complex (RSC), anterior cingulate and medial prefrontal cortex area 33 prime (33pr), piriform cortex (Pir), anterior agranular insula complex (AAIC), Entorhinal Cortex (EC), presubiculum (PreS) and hippocampus (H) as defined by the HCP-MMP1.0 parcellation^58^ based on the multi-modal atlas described by Glasser et al. (2016)^59^.

#### Myelin-informed source reconstruction visuo-motor paradigm

The resulting myelin-informed forward models were then used to solve the inverse problem for broadband visual oscillatory activity (0-80 Hz) within the time window of 0-200 ms following the onset of gratings. We restricted our analysis to trials where gratings were presented in the left hemifield after the 8 seconds inter-stimulus interval. In all source reconstructions, a data complexity reduction step was performed by projecting the MEG data onto 16 temporal modes. For the MSP approach, we used the default standard library of 256 patches with a patch width of 6 mm full-width of half maximum. As for the simulated data, we used the classic Bayesian source inversion scheme in SPM12 across source reconstruction approaches to enable a fair comparison of generative models.

## Results

### Single dipole simulations

We first perform single dipole simulations to test our fundamental assumption that increasing local leadfields to account for the expected increased current source density at sites of increased local myelin density can significantly improve source reconstruction accuracy. Hereby, we investigate the impact of the chosen leadfield scaling and SNRs on source reconstruction accuracy, evaluated by using DLEs and log model evidence with respect to a null model without leadfield scaling, across three popular source reconstruction approaches. Results are summarised in Figure 1. For the EBB and MSP approaches, DLEs tended to increase with decreasing SNR for the null model without any leadfield scaling as expected. Dipole localisation errors were highest for the MNE approach and remained relatively stable across the SNRs tested, while DLEs were comparatively small for the MSP approach, but showed a larger variability.

**Fig. 1.**
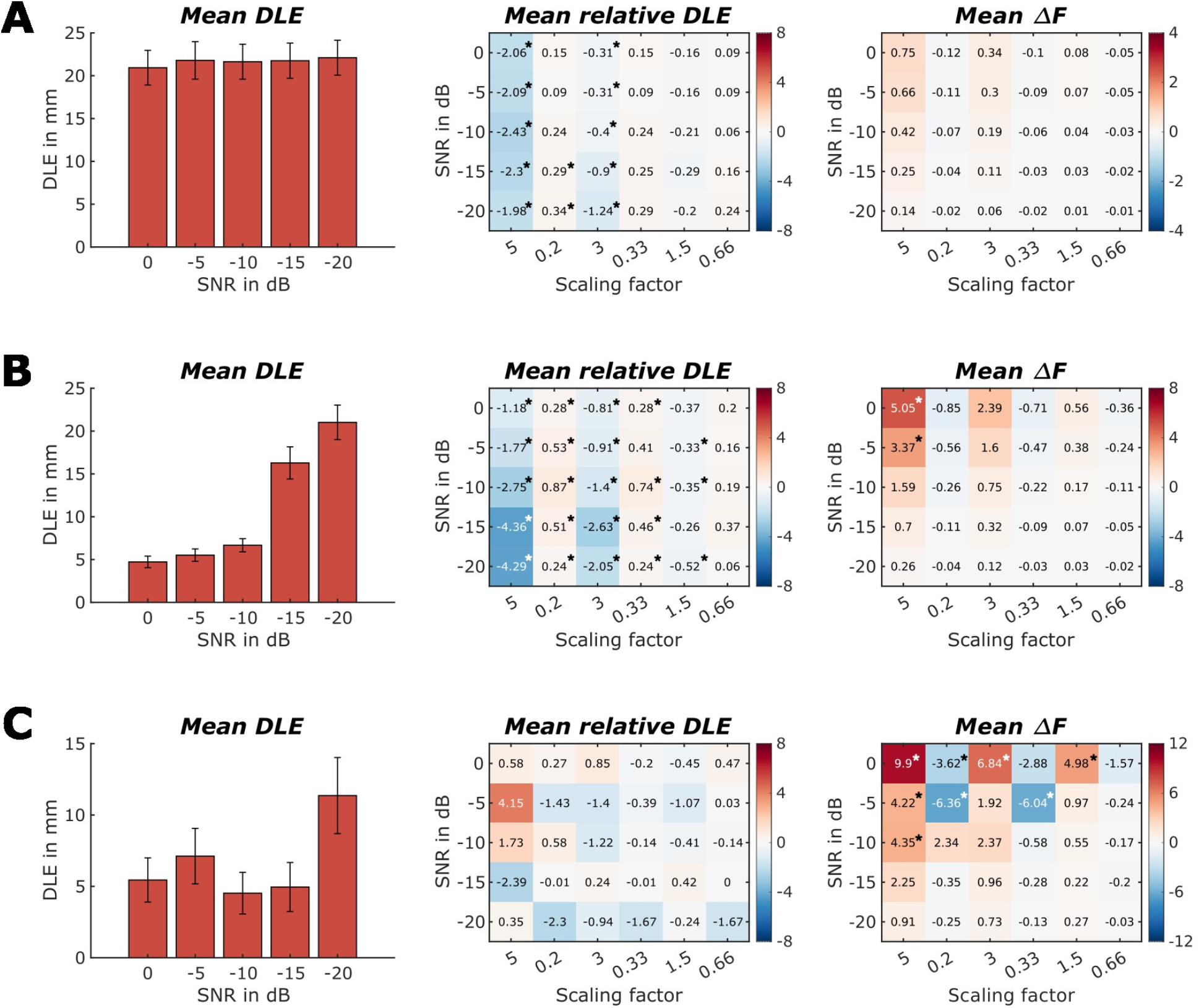
Dipole localisation errors (DLEs) and Bayesian log model evidence differences for locally increased or decreased myelin estimates. For **A** Minimum Norm Estimates, **B** Empirical Bayesian Beamformer and **C** Multiple Sparse Priors source reconstruction approaches. Left row panels show mean DLEs across signal-to-noise ratios for a null model without leadfield scaling. Middle row panels show relative DLEs in mm across varying leadfield scaling factors with either locally increased or decreased leadfields at the simulated cortical patch relative to the null model. Right row panels show log model evidence differences across SNRs and leadfield scaling parameters relative to the null model. Asterisks indicate significant differences from the null model for DLEs and Free Energy values. All results are based on 60 simulations per scaling factor and SNR for each source reconstruction approach.

DLEs decreased for simulations with locally increased leadfields at the simulation patch site and increased for locally diminished leadfields for the MNE and EBB approach (Fig. 1 A and B, middle column). Those differences were significant at scaling factors *f*_*s*_ of 5 and 3 for both MNE and EBB source reconstruction approaches and additionally at a smaller scaling factor of 1.5 for the EBB approach. We also found increases in DLE at scaling factors that resulted in decreased local leadfields (*f*_*s*_ = 0.2 for MNE and *f*_*s*_ = 0.2 and 0.33 for EBB). We did not find any significant differences between DLEs of myelin-informed forward model and the null model without leadfield scaling for the MSP approach, an observation we ascribe to the larger variance of DLEs when using multiple sparse priors in source reconstruction.

We find that the Bayesian model comparison framework is sensitive to changes in myelin for the MSP and EBB source reconstruction approaches at moderate-to-large sensor-level SNRs (≥10 dB) (Fig.1, right column). For the EBB approach, differentiating between myelin-informed forward models and the null model was only feasible at a scaling factor of 5. For the MSP source reconstruction approach, we were able to distinguish a myelin-informed forward model from the null model at a smaller scaling of 1.5. For the MNE approach, the relative log model evidence for the myelin-informed forward models was similarly positive for increases and negative for decreases in leadfield strength. However, these log model evidence differences were not significantly different from the null model across all scalings and SNRs tested.

We next tested whether the effect of myelin-informed leadfields is spatially specific (Fig. 2). Relative log model evidence for a control condition where we changed the myelination at a patch at the opposite hemisphere indicates that this is indeed the case: We did not observe any significant model evidence differences between the null model and myelin-informed control models. Minute systematic changes in log model evidence at larger myelin scalings for the MNE and EBB approaches (Fig. 2A, B) may be due to less source strength being needed to explain sensor noise which can be modelled using a current dipole at this patch.

**Fig. 2.**
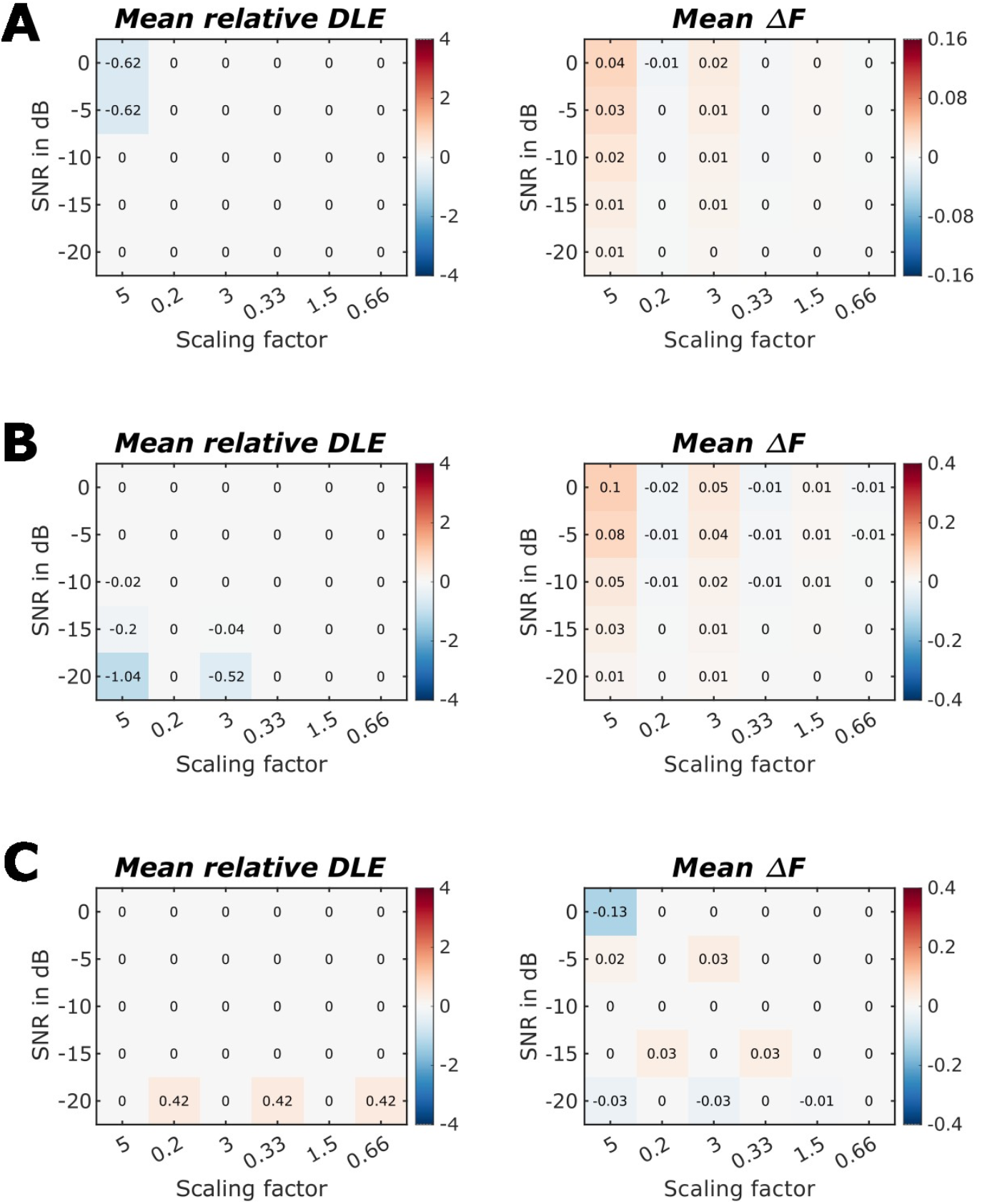
Relative dipole localisation errors (DLEs) and Bayesian log model evidence differences for locally increased or decreased leadfields at control sites at the opposite hemisphere. For **A** Minimum Norm Estimates, **B** Empirical Bayesian Beamformer and **C** Multiple Sparse Priors source reconstruction approaches. Left row panels show DLEs across varying leadfield scaling factors with either locally increased or decreased leadfields at a patch opposite to the cortical patch where the current dipole source was simulated. Right row panels show log model evidence differences across SNRs and leadfield scaling parameters for the same control models. No significant differences from the null model were observed for DLEs and Free Energy values. All results are based on 60 simulations per scaling factor and SNR for each source reconstruction approach.

We further examined the impact of leadfield strength at the adapted simulated source patch locations on our ability to distinguish myelin-informed forward models from the null model. This investigation was motivated by previous studies that demonstrated that discrimination accuracy is affected by leadfield strength^5,6^. We observed that differences in model evidence strongly vary with the leadfield norm at the peak vertices of simulated patches: At source locations with larger leadfields, and thus a strong impact on the sensor data, applying myelin-informed leadfield adaptation had a more pronounced effect than at locations where sources had a lesser impact on the sensor data (Fig. 3B, C). We found that a quadratic model explained this relationship significantly better than a linear fit 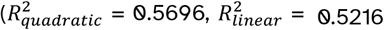; F-statistic: 11.5152, p <.01).

**Fig. 3.**
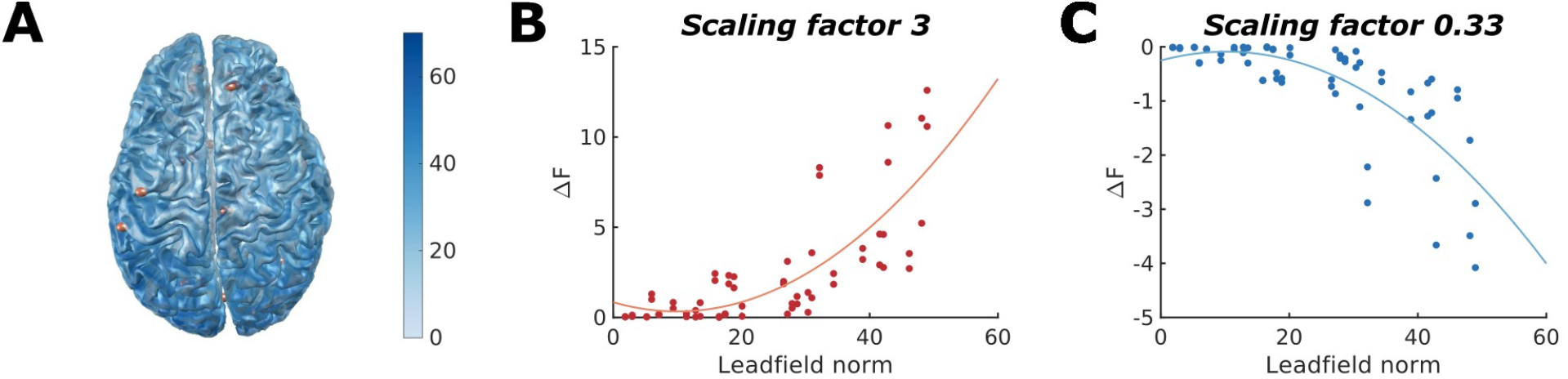
Impact of leadfield strength on the relative log model evidence differences for myelin-informed source reconstruction using the EBB approach at an SNR of 0 dB. **A** Shades of blue indicate the leadfield norm at each vertex across the mid cortical surface. Centre locations of the patches used as source locations are indicated by red spheres. **B** Correlation between peak vertex leadfield norm and log model evidence difference across the 60 simulation patches for a leadfield scaling of 110%. **C** The same correlation for a leadfield scaling of 91%. Fitting curves are based on a quadratic polynomial fit.

### The benefit of using myelin-informed leadfields decreases with increasing co-registration error

Next, we examined the impact of co-registration errors on our myelin-informed source reconstructions across a set of simulations with a leadfield scaling factor *f*_*s*_ of 5 at an SNR of 0 dB. We found that the benefit of using myelin-informed leadfields, as measured by the relative log model evidence, diminishes with increasing co-registration error for all tested source reconstruction approaches (Fig. 4). Differences in log model evidence did not significantly differentiate between myelin-informed and null forward models at co-registration errors exceeding 3 mm and 4 mm for the EBB and MSP source reconstruction methods, respectively. As previously shown in Fig. 1, for the MNE approach, differences in log model evidence from the null model were smaller than 3 and thus non-significant, even in the absence of co-registration errors.

**Fig. 4.**
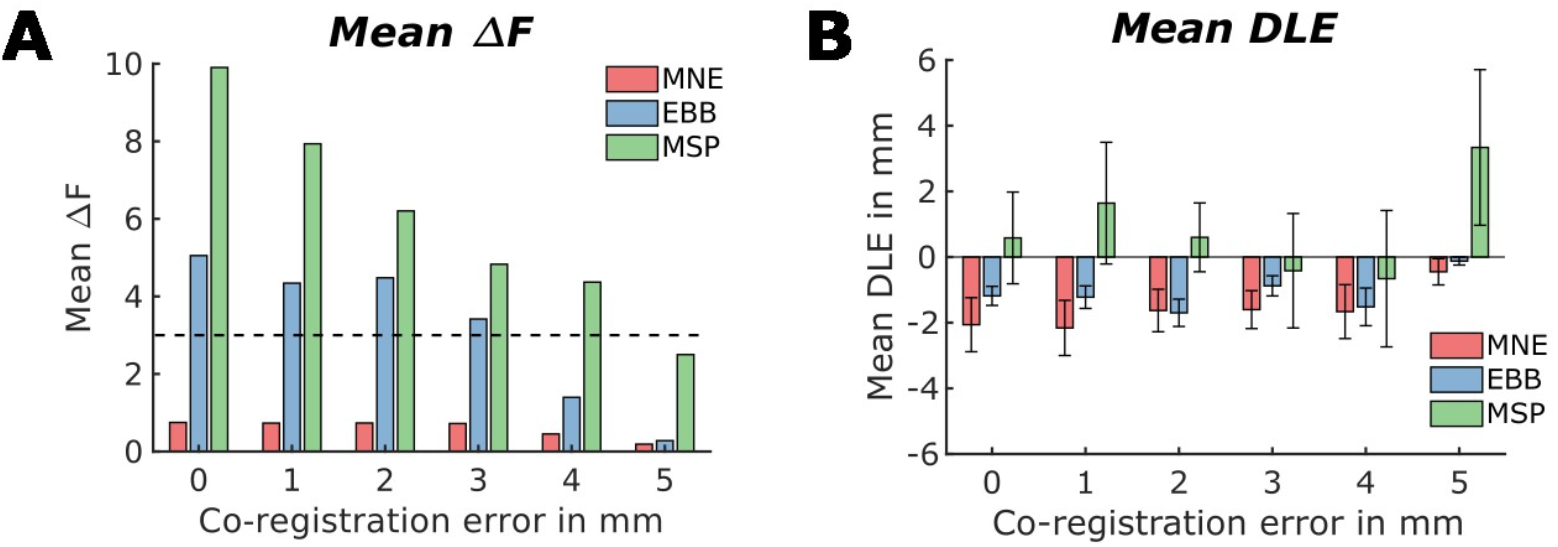
Impact of co-registration errors on the relative log model evidence differences for myelin-informed source reconstruction. Results are displayed for the EBB approach at an SNR of 0 dB and with a leadfield scaling factor of 5. **A** Mean relative log model evidence across co-registration errors for the three source reconstruction approaches tested. Log model evidence differences greater than 3 indicate that the myelin-informed forward model could significantly better explain the data than the null model without leadfield scaling **B** Mean relative dipole localisation errors across co-registration errors.

Paired t-tests showed that DLEs were significantly smaller than those for the null model at co-registration errors less than 5 mm for the MNE and EBB source reconstruction approaches. For the MSP approach, in contrast, DLEs did not differ significantly from the null model without leadfield scaling, regardless of the co-registration error applied.

### Inferring myelination patterns from MEG data using myelin-informed forward models

In a next step, we aimed to infer myelination patterns, consisting of two bilateral patches with opposite scaling directions, which underlie our simulated sensor data. Results are summarised in Fig. 5. Using the EBB source reconstruction approach, we found that employing a forward model with the same myelin scaling pattern as the one used to generate the sensor data increased the relative log model evidence compared to the null model at large scaling factors. This was observed for source reconstructions of simulated data generated using bilateral scaling patterns of 10 - 0.1 and 0.1 - 10.

**Fig. 5.**
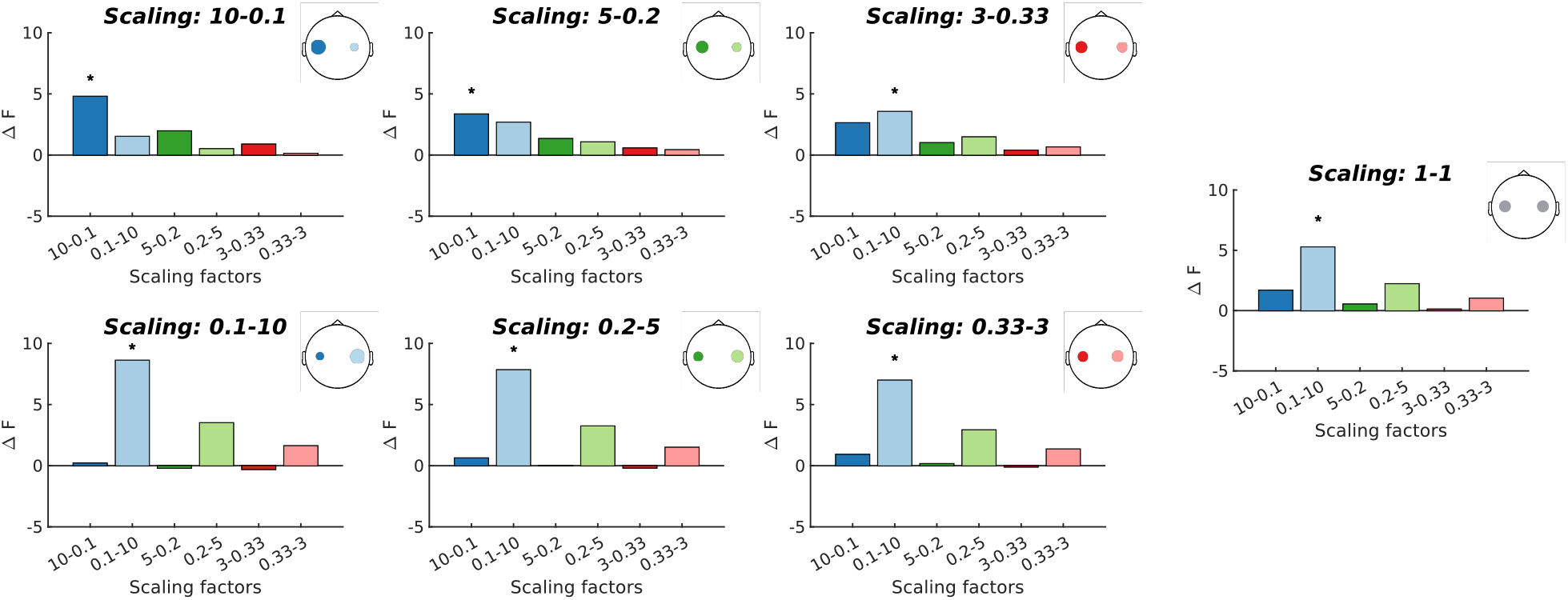
Inferring cortical myelination pattern using myelin-informed generative models. Each panel displays the relative log model evidence differences across various scaling patterns for sensor data generated by the myelination pattern indicated by the head inlays. Source reconstructions were conducted using the EBB approach at an SNR of 0 dB. Asterisks indicate log model evidence differences greater than 3 from the null model without leadfield scaling.

However, for weaker scaling patterns, we found that the largest log model evidence differences typically occurred at the highest scaling factors, regardless of the scaling strength used to generate the data. For example, when a pattern of scaling factors of 5 – 0.2 was used to generate the sensor data, applying the myelin-informed forward model with a more pronounced scaling pattern of 10 – 0.1 resulted in a larger model evidence difference than applying the myelination pattern 5 – 0.2 that was used to generate the sensor data. Similar results can be observed for the scaling pattern in the opposite direction, as shown in the bottom row of Fig. 5. Furthermore, for data generated using a smaller myelin scaling pattern of 3 – 0.33, the winning model was the scaling pattern 0.1 – 10, i.e., a myelination pattern in the opposite direction. We attribute this to a bias present already in the non-adapted forward model (scaling pattern 1 – 1, right-most panel in Fig. 5).

In summary, while we were able to successfully infer the directionality of the underlying myelination pattern in most cases, the strength of the applied scaling could not be inferred successfully by comparing the source reconstruction model evidence across different myelin-informed forward models. Additionally, biases present in the null model without adapted leadfields may interfere with our ability to correctly infer the underlying myelination pattern.

### Application to empirical MEG data from a visuo-motor experiment

Finally, we applied myelin-informed forward models to MEG data from a visuo-motor experiment and investigated how this affected the log model evidence for source-reconstructed visual activity in response to the grating onset for all three source reconstruction approaches tested. Fig. 6A displays the MNE source reconstructed oscillatory activity of a single participant using a subject-specific forward model without myelin-informed leadfield scaling. The broadband source-reconstructed activity, up to 80 Hz, shows a maximum at right early visual areas, as expected for gratings displayed at the left hemifield. Figure 6B shows the group-average myelin map derived from quantitative R1 maps, revealing higher myelination estimates at primary sensory and motor areas, in accordance with known cortical myeloarchitecture. To incorporate the myelin information into the source reconstruction of the experimental data, we scaled the leadfields in the forward model for each participant by the subject-specific normalised R1 values, multiplied by a set of scaling factors.

**Fig. 6.**
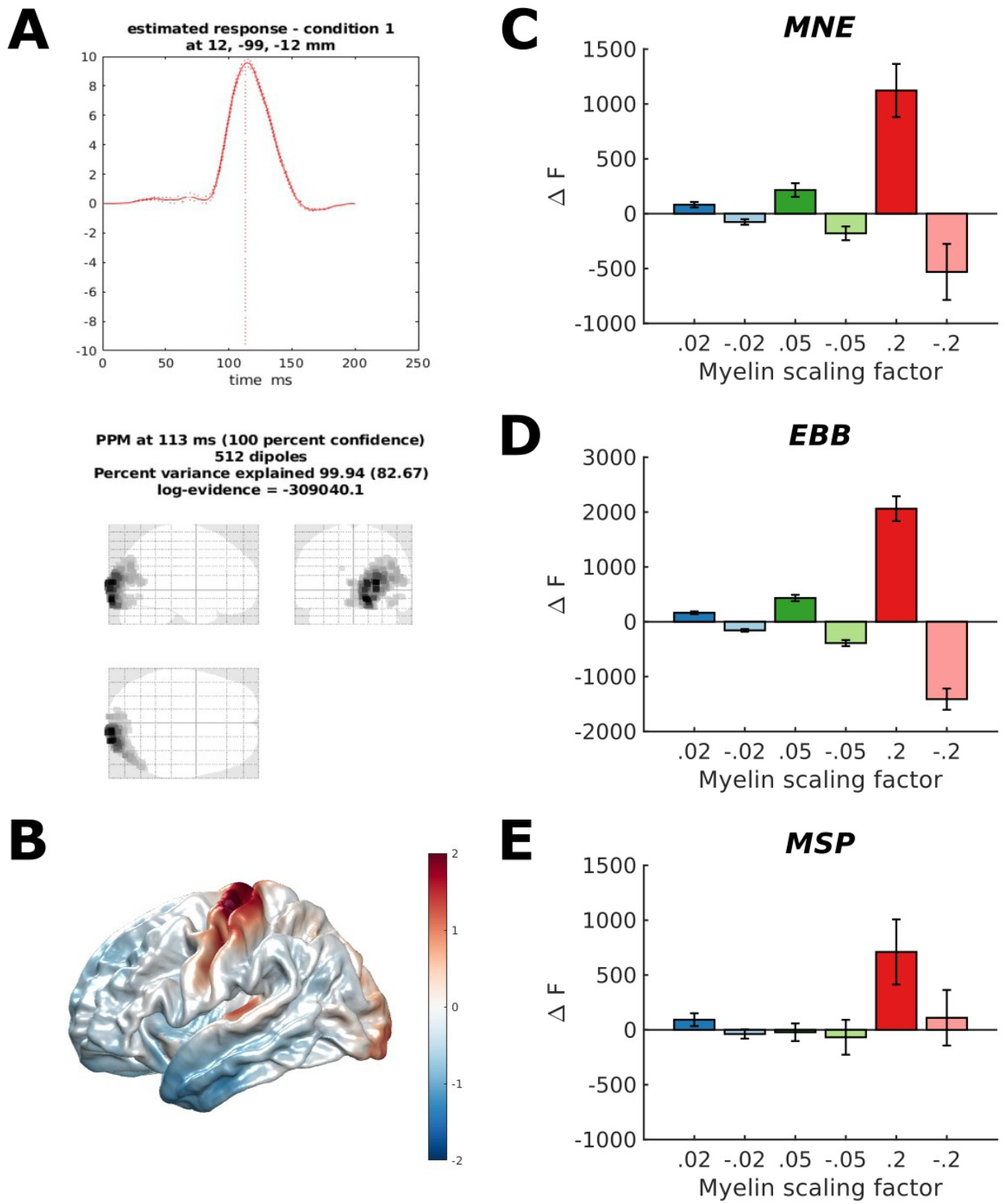
Myelin-informed source reconstruction of gradient-induced visual activity. **A** Broad-band MNE source reconstructed activity 0-200 ms after visual grating onsets for an exemplary participant. **B** Average normalised myelination across participants projected onto Freesurfer’s fsaverage mid surface. **C-E** Relative log model evidence across a set of positive and negative scaling factors for the three source reconstruction approaches.

For the MNE and EEB source reconstruction approaches, we find that adapting the forward model by scaling the leadfields with subject-specific myelin estimates based on the R1 maps and a positive scaling factor increased log model evidence compared to the null model (Fig. 6C-D). This increase in relative log model evidence grew with increasing scaling factors and continued to do so even at biologically implausible high myelin scaling factors (Suppl. Fig. S1). For negative scaling factors - which correspond to the hypothesis that increased myelination reflects a lower cortical current density - we observed a decrease in relative log model evidence, which again increased with decreasing scaling factors (Fig. 6C-D). We found this pattern of relative model evidence across positive and negative scalings to be consistent across subjects (for 12 and 13 out of 14 participants for the MNE and EBB source reconstruction approaches, respectively, as shown in supplementary Figs. S2 and S3).

For the MSP approach, the winning model was the forward model with the scaling factor 0.2 (Fig. 6E). However, this finding was not consistent across participants, where the same model was the winning model only for half of the participants (7 out of 14 participants, Fig. S4).

## Discussion

We demonstrate using simulated and empirical data that myelin-informed generative models can be used to improve M/EEG source reconstruction accuracy. Myelin-informed forward models with positive leadfield scalings at the location of the simulated cortical patch significantly improved source reconstruction accuracy at high and moderate SNRs. Further, we were able to infer the directionality of the scaling pattern for two dipole simulations. However, the model with the strongest scaling was generally the best model, irrespective of the magnitude of the original scaling. Additionally, biases present already in the null model further limit our ability to infer cortical myelination patterns from MEG data. For the visuo-motor MEG data, informing the forward model by subject-specific myelin estimates improved source reconstruction accuracy of visual responses. In the following, we discuss our results in more detail.

### Single dipole simulations

For the simulated data involving single dipole activity, we found that the model comparison framework is sensitive to changes in myelin in simulations at high to moderate sensor-level SNRs (> =10 dB) for the MSP and EBB source reconstruction approaches. We thus provide evidence that adapting leadfields to incorporate information about cortical myelination into forward models can significantly affect source reconstruction accuracy. Relative log model evidences were not significant for the MSP approach. We assume that the source covariance prior of the MNE approach is not well-suited for explaining the focal activations simulated in our study, which may compromise our ability to discriminate between different myelin-informed models.

For the EBB approach, we only found significant log model evidence differences for a scaling factor *f*_*s*_ of 5. As mentioned previously, a scaling parameter of 5 translates, on average, to an approximately 1.8-fold increase in leadfield strength at the peak vertex compared to the null model. Assuming a direct correspondence between leadfield adaptation and cell density, this implies that we can distinguish between cell density differences that conform to the known variability of cell density across cortical areas^60,61^, albeit they are relatively large. Note that for the MSP approach, we also found significant changes with respect to the null model for a forward model that reflects smaller changes in myelination and hence cell density (*f*_*s*_ = 1.5, corresponding to an approximately 10% increase in leadfield strength at the peak vertex).

The benefit of using myelin-informed forward models we observed for the EBB and MSP approaches was spatially specific. Forward models where the leadfields were adapted at locations opposite to those where the cortical activity was simulated did not differ significantly from the null model.

We found that adapting leadfields with a higher leadfield norm had a stronger impact on relative model evidence than those with a lower norm. While this correlation between leadfield norm and relative model evidence may seem trivial (as we scale the leadfields by a multiplicative factor), it highlights the fact that differences in myelination at sources close to the sensors will have a greater impact on the expected current density than myelin changes at deeper sources that are farther from the sensors.

#### Impact of co-registration errors

We found that the benefits of incorporating myelin-informed leadfields diminished with increasing co-registration errors, with no significant improvement over the null model at a co-registration error of 5 mm. As expected, the added uncertainty about head location from increased co-registration errors compromises our ability to discriminate between models. With increasing co-registration error, we are less able to explain the sensor data accurately and incorporating the myelin priors is rendered less efficient. Fortunately, we now have the tools to achieve high-precision measurements. In cryogenic MEG using subject-specific headcasts can provide the necessary co-registration accuracy^49,50^, while the use of rigid helmets^62^ combined with high-accuracy optical scanning devices are expected to provide sufficient co-registration accuracy in OPM-MEG systems^63,64^.

### Inferring myelination pattern from MEG data

We were able to infer the directionality, but not the scaling strength of the myelination patterns used for MEG data generation. We presume that this is because current dipole sources at locations of increased myelination (increased leadfield strength) have a stronger impact on the sensor level patterns than current dipole sources at locations of decreased myelination (decreased leadfield strength). Therefore, less source power is needed to explain the sensor pattern arising from the stronger dipole source, which introduces a bias towards explaining the dipole with the stronger impact (at the expense of modelling the weaker dipole). This reasoning can also explain the bias we observed for the null model: since the bilateral source locations were defined on a cortical surface derived from experimental data, dipole locations were not perfectly symmetric across the two hemisphere and could differ in the leadfield strengths. The resulting asymmetry in simulated current dipole strengths in turn may render it more favourable to use one of the scaled forward models than the null model for source reconstruction. We were able to further corroborate this explanation by showing a significant negative correlation between the difference in leadfield norm between bilateral patch pairs and the relative log model evidence differences for the forward model with the 0.1 – 10 scaling across simulations (Spearman correlation coefficient: -0.7935, p <.0.00001).

While we expected a cut-off where increasingly large scaling factors would again lead to a decrease in model evidence (as neglecting the weaker dipole entirely makes it increasingly more difficult to represent the sensor level pattern accurately), we found that even extreme and biologically implausible scaling pairs yielded large positive log model evidence differences. We speculate that this observation can be explained by the decrease in source power needed to explain the stronger current source surpassed the disadvantage of having to model the sensor activity due to the weaker current source, as the latter can still be approximated by neighbouring vertices, albeit at the cost of increased model complexity^65^.

Our ability to infer the true underlying myelin density by means of myelin-informed forward models and Bayesian model comparison is thus limited.

### Application to MEG data from a visuo-motor experiment

For the visuo-motor MEG data, we were able to demonstrate that informing the forward model using subject-specific myelin estimates can improve source reconstruction accuracy across participants when using the MNE and EBB source reconstruction approaches.

It is important to note that our approach is situated in a Bayesian framework and results need to be interpreted as such. Here we used MEG data from a visuo-motor task where the oscillatory responses to visual gratings are known to be generated in primary visual areas^66^. Myelin estimates from the R1 maps showed the expected pattern of more strongly myelinated primary sensory and motor areas and less heavily myelinated association areas^67,68^, which means that the leadfields corresponding to the primary visual areas were increased in the myelin-informed forward models for positive scalings. This is expected to result in an advantage of the myelin-informed over the null model. However, if our estimated source activity would have originated from a less myelinated area, using such a myelin-informed forward model would have been disadvantageous compared to the null model without leadfield scaling. Consequently, if we anticipate sources to originate from less myelinated regions, this knowledge should be incorporated as an additional prior^42,69^.

Using the MSP approach, we were unable to differentiate between the different myelin-informed forward models. In contrast to the simulated data, for the experimental data, patch centres do not necessarily match the current source locations. This reduced ability to accurately represent the experimental data may have compromised our capability to distinguish between myelin-informed forward models. The higher variability in relative model evidence due to the higher number of hyperparameters that need to be estimated for source inversions with MSP priors may also have contributed to the inconsistent results.

We note that the empirical data was acquired using headcasts, with improved SNR and reduced co-registration errors in comparison to conventional MEG, and thus represent data from favourable measurement conditions. Other studies aiming to apply myelin-informed forward models in MEG source reconstruction, should be aware of the requirements on SNR and co-registration errors when acquiring the data.

With a data acquisition time of below half an hour, MPMs can be easily acquired next to M/EEG data, where they can also be used instead of a standard T1w structural scan to set up the forward model^70^. Furthermore, advances in ultra-high-field qMRI techniques, with resolutions reaching 500 μm or higher, hold great promise for extending our approach to laminar-specific myelin-informed models^5,7^ and may enhance the reliability of inferring the laminar origin of MEG signals.

### Limitations

There are several caveats and limitations to this study. Although existing studies have established a link between neural activity and cortical myelination^29,30,71^, and our previous work has demonstrated that cortical myelin density estimates from qMRI predict the strength of neurophysiological responses^38^, direct empirical evidence confirming that cortical current density varies systematically across the cortex in accordance with myeloarchitecture is still lacking. We would also like to highlight that Murakami and Okada (2015)^72^ emphasise the invariance in maximal current dipole moment density across a wide range of brain structures and species. Nevertheless, the range of current densities they report across human neocortex - from 0.16–0.77 nAm/mm^2^, representing nearly a five-fold difference between the minimum and maximum observed values - is substantial enough to account for the variability in myelin density across cortical areas that we have assumed in our study.

It is also possible that myeloarchitecture is more closely associated with brain connectivity^36,71,73^ or the temporal and spectral features of neurophysiological activity rather than with dipole moment strength. For example, Hunt et al. (2016) identified a significant correlation between the structural covariance of cortical myelination and electrophysiological networks of neural oscillatory activity. In another study, Shafiei et al. (2023)^74^ found that the dominant spatial gradient of neurophysiological dynamics reflects characteristics of power spectrum density and linear correlation structure of the signal, and covaries with several micro-architectural features, including the cortical myelination gradient from early sensory and motor areas to associative areas. Relatedly, Mahjoory et al., 2020^75^ reported in resting-state MEG that the dominant peak frequency in a brain area decreases significantly along the cortical hierarchy.

Assuming for now that cortical myelin density is predictive of MEG signal strength, we must further appreciate that R1 maps (like other qMRI measures) can only serve as a proxy for myelin content. Quantitative MRI parameter maps are sensitive to multiple tissue components to a varying extent^76^ and thus lack the specificity to directly infer the abundance of any single microstructural tissue component.

In the present study, we assumed a linear relationship between myelin estimates from quantitative MRI and the strength of MEG signals. Future research could investigate more complex models that link qMRI-based myelin estimates and functional MEG or EEG signals. Validating these models by comparing their ability to explain experimentally measured data may provide valuable insights into microstructure-function relationships in the living human brain. However, our observation that we were only able to infer the directionality, and not the strength, of a simple bilateral myelination pattern in our simulations warrants caution. Finally, we note that microstructural and functional gradients have been reported to be increasingly dissociated in transmodal cortices^77^ and that the link between myelination and MEG measures may thus vary across cortical regions, adding an additional layer of complexity.

## Supporting information

Supplemental results

## Acknowledgements

The authors gratefully acknowledge funding from the European Research Council under the European Union’s 23 Seventh Framework Programme (FP7/2007-2013)/ERC grant agreement no 616905 and the Medical Research Council UK MEG Partnership grant MR/K005464/1. SH was supported by a scholarship from the Pontius foundation. The Wellcome Trust Centre for Neuroimaging is supported by core funding from the Wellcome Trust 091593/Z/10/Z.

