## Supplemental results for "Myelin-informed forward models for M/EEG source reconstruction"

### Supplementary material

#### Averaged log model evidence for the myelin-informed source reconstructed visual activity at extreme scaling factors

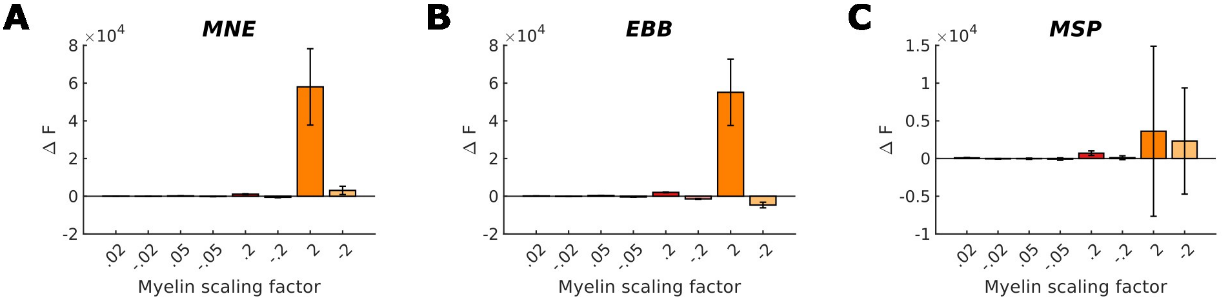


**Fig. S1** Relative log model evidence across a set of positive and negative scaling factors for the three source reconstruction approaches. Asterisks indicate significant differences from the null model without any leadfield scaling.

#### Relative log model evidence for the myelin-informed source reconstructed visual activity across participants

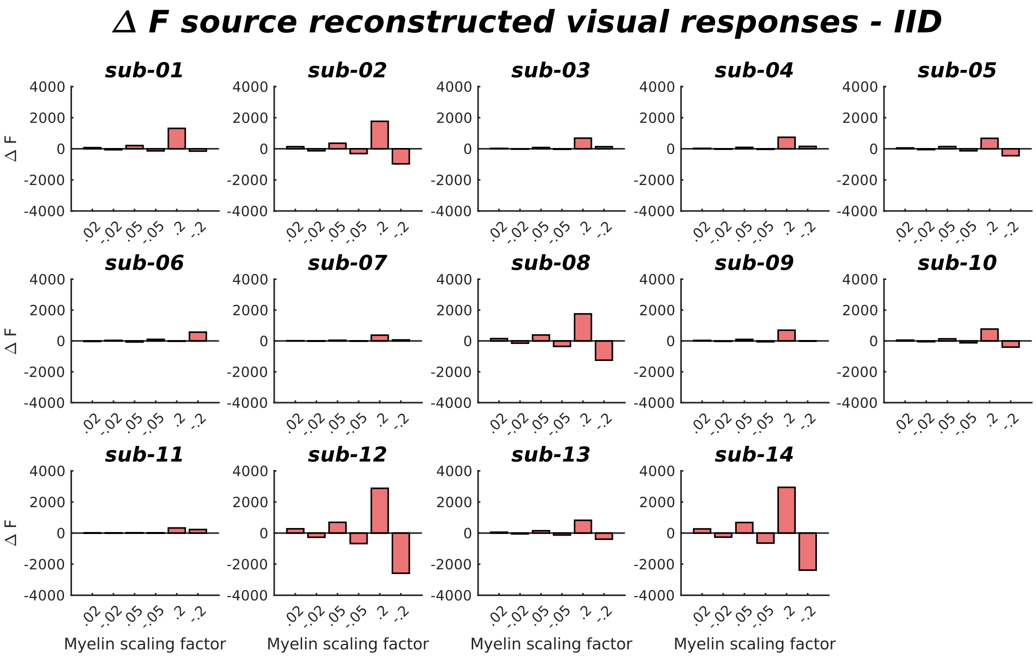


**Fig. S2 Relative log model evidence for myelin-informed source reconstruction of broad-band activity 0-200 ms after visual grating onsets across participants for the MNE source reconstruction approach.** Normalised myelin estimates derived from qMRI were multiplied with a scaling factor and used to scale the leadfields. Relative log model evidence was assessed across a set of positive and negative scaling factors.

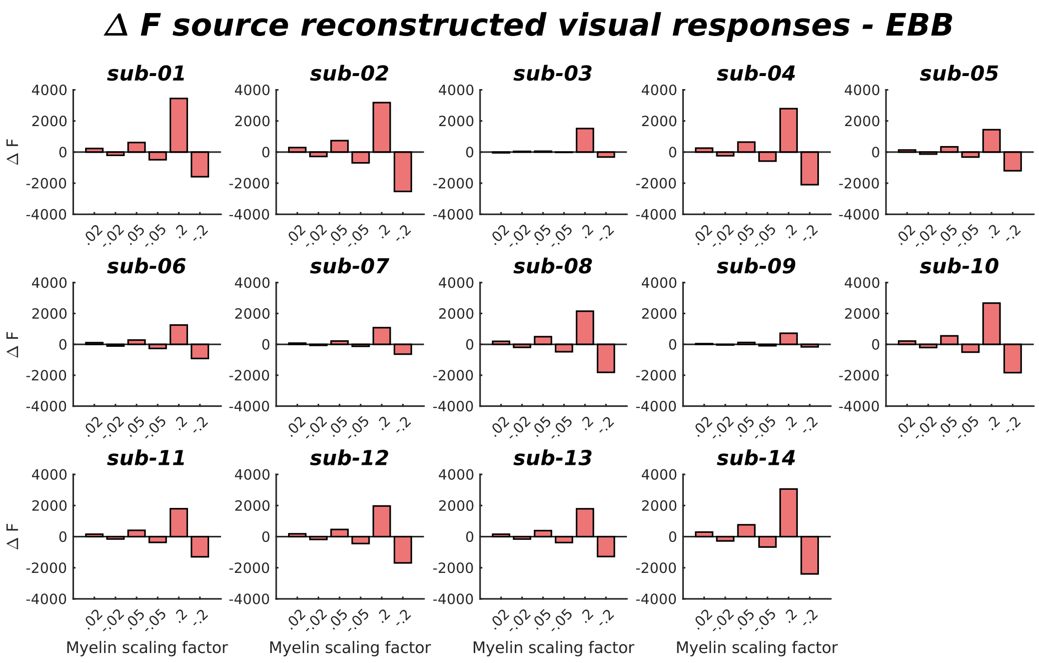


**Fig. S3 Relative log model evidence for myelin-informed source reconstruction of broad-band activity 0-200 ms after visual grating onsets across participants for the EBB source reconstruction approach.**

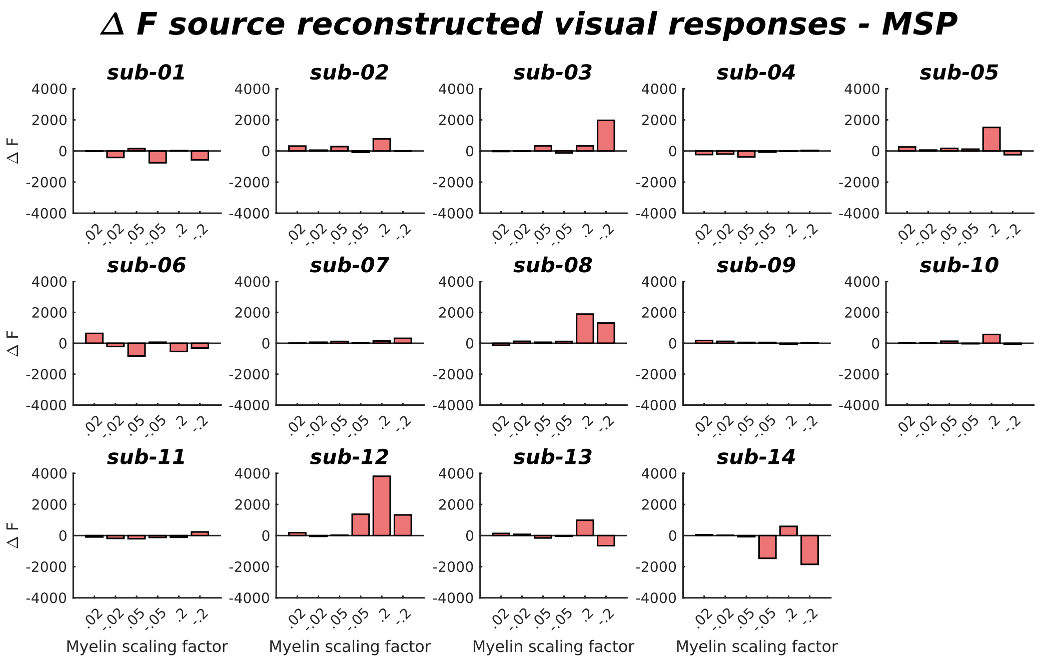


**Fig. S4 Relative log model evidence for myelin-informed source reconstruction of broad-band activity 0-200 ms after visual grating onsets across participants for the MSP source reconstruction approach.**
